# Migration strategy varies with novel environment response in common noctule bats

**DOI:** 10.1101/2022.12.22.521583

**Authors:** Theresa Schabacker, Sofia Rizzi, Tobias Teige, Uwe Hoffmeister, Christian C. Voigt, Lysanne Snijders

**Affiliations:** Department of Evolutionary Ecology, Leibniz Institute for Zoo and Wildlife Research, Berlin, Germany; Freie Universität Berlin, Berlin Germany; Natural History Museum, Leibniz-Institute for Evolution and Biodiversity Science, Berlin, Germany; Humboldt Universität zu Berlin, Berlin, Germany; Büro für faunistisch-ökologische Fachgutachten, Berlin, Germany; natura – Büro für zoologische und botanische Fachgutachten, Germany; Behavioural Ecology Group, Wageningen University, Netherlands

**Author notes:** Current affiliation of corresponding author: Natural History Museum, Leibniz-Institute for Evolution and Biodiversity Science, Berlin, Germany Contact.

**Keywords:** animal migration, bats, echolocation, exploration, novel environment, stable isotope analysis

## Abstract

Global ecosystems are changing dramatically due to land transformation and climate change. Global change is a particular challenge for migratory animals that rely on multiple stepping stones on their journeys. Migratory animals have a range of strategies to accomplish this, but not all of these strategies may be appropriate for the challenges ahead. Understanding the variation in migratory strategies and their behavioural correlates is therefore critical to understand how vulnerable species will be in the future, especially in endangered and elusive taxa such as bats. Here, we combined isotopic geolocation with an in-situ behavioural assay to investigate whether behavioural responses to a roost-like novel environment correlated with variation in migration strategies (local or distant origin based on isotopic geographic assignments), in the partially migratory bat, *Nyctalus noctula*. We quantified emergence behaviour, spatial activity, and echolocation call activity. Local bats were more likely to emerge into the novel environment than bats from more distant origins. However, local and distant bats did not differ in spatial activity and acoustic exploration (relative call activity per space unit). Our findings indicate that local bats may more pro-actively cope with novelty, but that acoustic exploration is equally important for local and migratory bats during explorations.

## Introduction

Animal migration is thought to mitigate the effect of seasonal changes in environmental conditions on animals, thereby increasing survival and fitness (Heape, 1931; Shaw, 2016; Shaw et al., 2012). A large number of organisms undertake migrations every year. In doing so, migratory animals depend on several different habitats and stepping stones on route, making them susceptible to changes in these environments. Changes in the environment are currently primarily caused by anthropogenic actions and the construction of human infrastructure, as wind energy facilities, roads or power lines, pose great threats to migratory populations. Migratory species suffer from direct (i.e., collision fatalities) and indirect (i.e., habitat fragmentation, migratory corridor disruption) impacts and are thus especially vulnerable (Hovick et al. 2014). Prior differences in selection pressures have led to a variety of migration strategies, both across species and within species (Dale et al., 2019). The adaptive value of these migration strategies may be changing in modern times due to anthropogenic impacts. Depending on their migratory strategy, animals may or may not be able to cope with the rapid changes currently been seen on our planet (Wilcove & Wikelski, 2008). Therefore, understanding the variation of migration strategies in context to environmental cues seems key for predicting how vulnerable migratory species are.

Partial migratory populations may offer interesting insights into the adaptive value of migratory strategies. Populations of partial migratory species comprise of individuals that remain local throughout the year and individuals that migrate over long distances. Partial migration is ubiquitous in various taxa, including invertebrates (Menz et al., 2019), amphibians (Grayson et al., 2011), fish (Espinoza et al., 2016), birds (Arnekleiv et al., 2022) and mammals (Purdon et al., 2018). In partial migratory populations, individuals of the same species respond with different migration strategies to the same environmental cues. For instance, migration strategies covaried with latency to emerge into a new environment in fish (Chapman et al., 2011) and the readiness to approach a novel object in birds (Nilsson et al., 2010). Yet, no study investigated if migrants and locals of the same species and population differ in how they explore, i.e. examine and investigate, their environment, a behaviour which is crucial for detecting novel resources and risks. Knowing whether migrants and locals vary in how they cope with novelty may help in predicting how migrants and locals differ in their vulnerability to human-induced environmental changes.

Despite their species richness, global abundance and pivotal role in ecosystem functioning (Ghanem & Voigt, 2012; Kunz & Fenton, 2005), little is known about migration in bats (Popa-Lisseanu & Voigt, 2009). Even bats of small size may travel several thousand kilometres during their seasonal journeys (Pētersons, 2004). Bats use energy-saving strategies during migration by travelling at optimal flight speeds (Troxell et al., 2019) and by entering torpor during stopover periods (McGuire et al., 2014). Wing morphology and immunological parameters of North American partial migratory bats differ between migratory and non-migratory conspecifics (Rogers et al., 2022). However, climate change poses new threats to migratory bats because they are exposed to novel human-made infrastructures, such as wind energy facilities, with cause high numbers of casualties in bats (Voigt et al., 2022). In a recent study, it was shown that mostly juvenile bats collide at and die at wind turbines, which might be related to their exploratory behaviour toward and naïve interaction with novel environmental cues such as wind turbines (Kruszynski et al., 2022). These findings stress the importance of individual behaviour towards novelty on the susceptibility to anthropogenic changes. Thus, as a first step toward an improved understanding of bat migration strategies, we examined the behavioural responses to a novel environment of a partially migratory bat, the common noctule bat, *Nyctalus noctula*. Based on isotopic geographic assignments, Lehnert et al. (2018) revealed a high variability in migratory strategies in European common noctule bats, providing evidence for differential and partial, female-biased migration in this species. Furthermore, they observed a high degree of consistency in migratory behaviour within individuals of common noctule bats. While morphological and physiological trade-offs for partially and differentially migrating bats have been described (Rogers et al., 2022), so far, we still lack evidence whether behavioural variation, specifically in response toward novel environmental cues, are linked to intra-specific migratory strategies in bats.

To investigate whether migration strategy and exploratory behaviour vary between individuals of a partial migratory species, we determined the migration strategies and quantified the emergence and subsequent exploration behaviour of 89 female common noctule bats. To this end, we used a novel approach combining isotopic geolocation with an in-situ maze-like novel environment assay in a partial migratory common noctule bat population. Bats offer a unique model to investigate exploratory behaviour as we can reliably quantify how they sample their environment via echolocation. This novel environment assay was recently established to quantify exploratory behaviour in another tree-dwelling bat species, the Nathusius’ bat, *Pipistrellus nathusii*, and revealed individually consistent differences in spatial activity and ‘acoustic exploration’, i.e., the level of environmental cue sampling per unit space (Schabacker et al., 2021). The experimental arena that was used in our behavioural assay mimicked a potential novel roost and we thus regarded its exploration as relevant for a tree-dwelling bat species. We differentiated between migratory and non-migratory strategies based on isotopic geographic assignments. This approach uses naturally occurring differences in stable isotope ratios, for example in stable hydrogen isotope ratios of precipitation water, which are governed by global-scale hydrologic processes. The isotopic differences in precipitation are seasonally and spatially predictable by latitude and elevation, and allow the reconstruction of so-called isotopic landscapes, i.e., isoscapes (Bowen et al., 2005; Courtiol et al., 2018; Hobson, 1999, 2008). This local isotopic signature is incorporated into the tissue of consumers via the diet and drinking water, and then conserved over specific time periods, depending on the metabolic (in)activity of tissues. Analysing stable hydrogen isotope ratios of keratinous material, which is a metabolically inert body product, reflects the stable hydrogen isotope ratios of the location where fur was grown (Fraser et al., 2013). Accordingly, the analysis of stable hydrogen isotope ratios allows the tracking of animal movements over geographic distances (Baerwald et al., 2008; Hobson & Wassenaar, 1996; Popa-Lisseanu et al., 2012; Sullivan et al., 2012; Voigt et al., 2014). We hypothesized that migratory strategies (local or distant geographic origin) of common noctule bats should correlate with their exploratory behaviour in our behavioural assay. We predicted migratory individuals to be more neophilic and exploratory than local conspecifics and therefore to emerge quicker and exhibit more spatial activity and acoustic exploration in a novel environment.

## Methods

### General fieldwork

Between April 11^th^ and 25^th^, and October 3^rd^ and 28^th^ of 2019, we sampled a total of 89 adult females from a wild colony of Noctule bats near the village of Prieros, Brandenburg, Germany (52°13’24”N, 13°45’16”E). We collected adult female bats in the early evening from roost boxes located 4-5 m above the ground. To minimize disturbance on the population, we selected roost boxes based on the presence of previously tested bats (identified by passive integrated transponders (PIT) tags, see below) or based on group size. Following removal from roost boxes, we determined sex, age, body mass (digital balance and spring balance) and forearm length (manual calliper). We excluded males from further study due to evidence of female biased migration in this species (Lehnert et al., 2018). Female bats were kept in groups in a dark environment (darkened plastic boxes, approximately 30 cm × 20 cm × 15) according to their association in the roost boxes where they were found. The holding boxes were equipped with cloth and heat packs to stimulate normothermia. Successively, we removed females from the boxes and conducted the behavioural assay on individual bats, primarily after sunset (N = 94) or just briefly before (N = 4; 35 to 9 minutes before sunset); a time during which common noctule bats are naturally active. Assays took place in a tent, within 200 m of the place of capture, providing a standardized environment to minimize external influences. Ambient temperatures were similar between the months with a mean of 10.9°C (range: 5.6 - 17.6°C) in April and 10.4°C (range: 6.0 - 18.0°C) in October. After the experiment we took a dorsal fur sample from just above the uropatagium for stable isotope analysis (Voigt et al., 2014). Dried fur samples were stored at ambient temperature until further analysis. Prior to release, we injected a PIT tag (Biomark, APT12, 12.5 mm, 134.2 kHz ISO FDX-B) subcutaneously in the dorsal neck region using an implanter (Biomark, MK25). PIT tags provided each individual with a unique ID-number which was readable with a mobile reader (Biomark, HPR Lite) at a distance of up to 25 cm from outside a roost box. We applied this method to facilitate the non-invasive localization of previously tested bats, yet, over both seasons, only 9 individuals could be recaptured and retested. Our assays included 89 unique female individuals (mean ± standard deviation: weight: 29.3 ± 4.1 g, forearm length: 54.2 ± 1.2 mm). All individuals were released at their respective roost boxes after testing. All fieldwork and associated procedures were conducted in accordance with the German law under a German animal welfare permit (number 2347-25-2018) and conservation permit (number LFU-N1-4743/128+25#314731/2018).

### Behavioural assay

Prior to an assay, we confirmed the normothermic condition of bats by measuring the skin temperature with a thermocouple (Peakmeter, PM6501; Thermocouple, Sensor SSP-1-150, Peakmeter, Shenzhen, China). Skin temperature (> 30 °C) is a non-invasive proxy for core body temperature (Barclay et al., 1996). In general, the behavioural assay simulated a novel tree roost with several chambers. Specifically, the maze-like arena (**Figure 1**; 40 × 40 × 5cm) consisted of nine separate chambers connected through small gates (3 × 2.5 cm) on the upper half of the walls. A rubber non-slip mat overlaid the floor to provide a good texture for crawling, a behaviour intuitive to tree-dwelling bats. A layer of insect screen overlaid the entire maze, preventing a bat’s escape and offering another climbing opportunity. The entrance to the maze was an opaque start tube (10 × 3cm), attached to the maze but blocked by a small wooden barrier, which was removed at the start of an assay. The entire maze was placed horizontally - to stimulate natural exploration behaviour in all directions - in a larger box (70 × 45 × 8 cm) with a transparent lid. We monitored the bat’s movement from a top-down perspective, by mounting a night vision camera (Sony Digital Camcorder, DCR-SR72E, Sony, Tokyo, Japan) on a tripod equipped with a horizontal arm, positioned 1.5m above the maze. Light was provided by an infra-red flashlight (T38, Evolva Future Technology, Shenzhen, China), shining at an angle from a fixed position. We recorded vocalizations with a directional USG Electret Ultrasound microphone (polar pattern 180°, Avisoft Bioacoustics/Knowles FG, Berlin, Germany) connected to an ultrasound recorder (UltraSoundGate 116Hb, Avisoft Bioacoustics, Berlin, Germany), which was placed within the larger box pointing to the centre of the maze.

**Figure 1.**
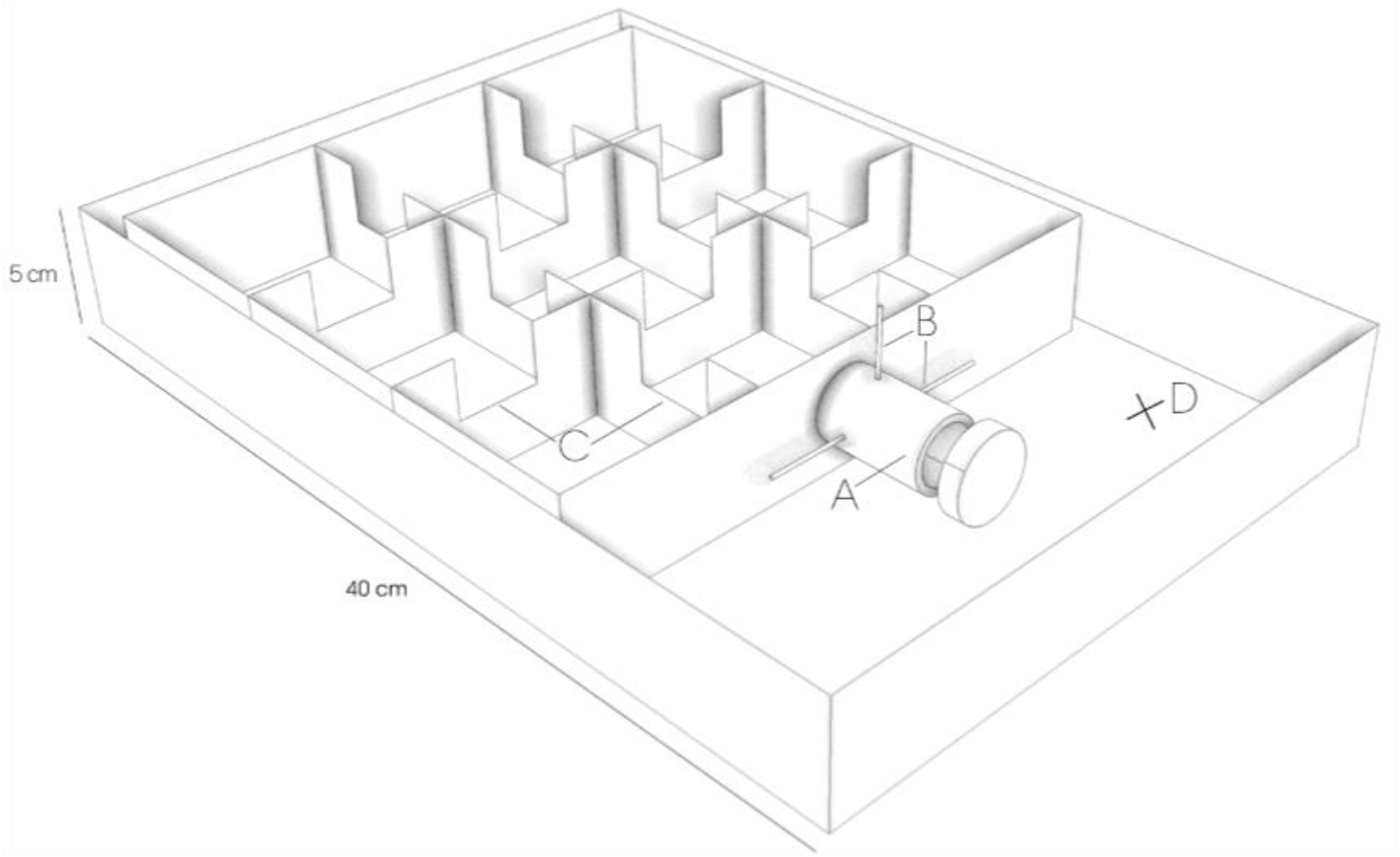
Schematic drawing of the maze used during behavioral assay. A) Opaque start tube where bats were placed at the start of each assay B) Barrier closing entrance to maze C) Gates connecting single rooms D) Position of microphone. ©Rebecca Scheibke

At the start of an assay, we placed the bat in the start tube, while a vertical barrier in the form of a small wooden dowel was still obstructing the exit. After an acclimatization period of 20 s, we removed the barrier and gave a maximum of three minutes for the bat to emerge. We recorded the bats’ emergence times, i.e., the time it took for an individual to emerge completely from the start tube. If a bat did not emerge during this time, we terminated the test. After a successful emergence, bats were allowed to explore the maze for two minutes. After each test, we removed potential olfactory cues by sterilizing the maze with a mild unscented detergent. Halfway through the night we rotated the setup to control for potential orientation biases. There was no significant correlation with the likelihood to emerge and the orientation of the setup (p > 0.3). We released the bats the same night, within 100 m of the original capture location.

### Stable isotope analyses

All stable hydrogen, nitrogen and carbon isotopes were analysed at the stable isotope lab at the Leibniz Institute for Zoo and Wildlife Research (Berlin, Germany). To prepare the samples for analysis, we cleaned off all external contaminants and oils with a chloroform:methanol (2:1) solution. To this end, samples were placed in the cleaning solution for 24 hours on a shaking platform (Phoenix GmbH). The fur samples were then dried for ten days in a drying oven (Heraeus Function Line, ThermoFischer Scientific, Bremen, Germany) at 50°C.

For stable hydrogen isotope analysis, we analysed samples in sequence with keratin reference materials with known stable isotope ratios for the non-exchangeable portion of hydrogen. We used four in-house standards: sheep wool from Sweden (−167.9 ± 1 ‰), sheep wool from Spain (−108.3 ± 1 ‰), goat wool from Tanzania, Africa (−66.3 ± 0.9 ‰) and USGS42 standard (−74 ± 0.5 ‰) from Tibetan human hair. A detailed description of the preparation of the standards is reported in Popa-Lisseanu et al. (2012). We weighed samples and standards using a microbalance (Sartorius ME5, Göttingen, Germany) to 0.274 ± 0.01 mg and placed them into 3.3 mm × 5 mm silver-foil capsules (IVA Analysetechnik e.K. Meerbusch, Germany). We folded capsules into small packages and stored them in a 96-well microtiter plate. After preparation of the plate, we left it in a drying oven at 50°C for 24 hours to remove extra moisture. Subsequently, we transferred samples and standards into the carrousel of a Zero Blank autosampler (Costech Analytical Technologies Inc., Italy) of the high temperature elemental analyser (HTO Elementaranalysator HEKAtech GmbH, Wegberg, Germany). The samples were flushed for one hour with chemically pure helium (Linde, Leuna, Germany) as a carrier gas at a flow rate of 100 ml/min, eliminating remaining moisture and suppressing the influx of ambient O_2_. The foil packages were then pyrolyzed in the element analyser at 1450°C. The gas chromatograph separates H_2_, N_2_ and CO at 80°C and introduces the resolving H_2_ through the Conflo III interface (ThermoFischer Scientific, Bremen, Germany) into the isotope ratio mass spectrometer (Delta V Advantage IRMS, ThermoFischer Scientific, Bremen, Germany). We used only those samples for further analysis where the amplitude peak of the δ2H sample did not exceed 6500 mV. This is based on repeated measures of in-house keratin standards better than 3‰ (which is equivalent to one SD). Consequently, we had to exclude eight samples (seven associated with first behavioural assays and one with a repeated assay).

For the analysis of stable carbon and nitrogen isotopes, we weighed samples to 0.35 ± 0.05 mg and placed them into 4 mm × 6 mm tin capsules (IVA Analysetechnik e.K. Meerbusch, Germany). The Flash EA 1112 Series Element Analyzer (Thermo Italy, Rhodano, Italy) combusted the fur samples and the Delta V Advantage isotope ratio mass spectrometer (Thermo Finnigan, Bremen, Germany), operating in continuous flow mode, measured the carbon and nitrogen stable isotope ratios of the combusted materials. Stable isotope ratios are reported in the delta notation relative to V-PDB (carbon) or AIR-N_2_ (nitrogen) scales as units per mill (‰). Laboratory standards (tyrosine and leucine) were calibrated as described in Popa-Lisseanu et al. (2012).

### Video analysis

We used the open-source event logging software BORIS (Friard & Gamba, 2016; version 7.9.1) for a detailed quantification of the behavioural responses. The response measures assessed here are congruent with those reported in Schabacker et al. (2021): (1) latency to head emergence (s), (2) latency to full body emergence (s), (3) duration of emergence (s), (4) the number of unique chambers discovered, (5) the total number of chambers visited and (6) number of times a bat poked its head in an adjacent chamber.

### Audio analysis

We recorded echolocation activity during the exploration of the novel environment and assessed the number of echolocation calls in Avisoft SASLab Pro (Avisoft Bioacoustics, Version 5.2). Preceding the analysis, we converted sampling frequency from 250 kHz to 150 kHz. Spectrogram computation was accomplished via Fast Fourier Transformation 256, parameters set to Hamming window (bandwidth 1270 Hz, resolution 977 Hz) and volume normalized to 75%. Vocal start and stop commands given by the observer allowed the synchronization of video and audio recordings. We identified distinct echolocation calls via Avisoft’s call detection and template-based spectrogram comparison feature, which we verified and corrected via visual inspection. In addition, we quantified the number of clearly distinguishable air puffs emitted by the noctule bats. The acoustic analysis thus resulted in two additional response measures: (7) total number of echolocation calls after full body emergence and (8) the number of air puffs. The number of air puffs was recorded for exploratory analysis, since we did not have a specific prediction for this response measure. Lastly, we quantified (9) ‘acoustic exploration’ as response measure, representing acoustic sampling of the environment using echolocation while accounting for spatial activity (number of chambers visited). Following Schabacker et al. (2021), this was calculated as the residuals of the total number of echolocation calls emitted over the total number of chambers visited during the assay. These residuals represent the variance that is not explained by the fitted regression line (see Results).

### Statistical analysis

All analyses were conducted using the statistical software R (R Core Team, 2022; version 4.2.1).

#### Isotopic geographic assignments

To delineate the origin of captured noctule bats in Brandenburg, we performed isotopic geolocation based on the R package “*IsoriX”* and tightly followed the workflow described in Courtiol et al. (2018). Briefly, the workflow is divided into three main components: generation of the isoscape, fitting the calibration model and geographically assigning unknown samples to the most probable place of origin. We built the isoscape based on hydrogen isotope ratios in precipitation water (δ^2^H_P_) obtained from the Global Networks of Isotopes in Precipitation (GNIP) database, which is a publicly available database of global isotope data. We fitted a geostatistical mixed model to predict the spatial distribution of δ^2^H_P_ roughly covering Europe (longitude range: −30° East to 60° West, latitude range: 30 North° to 70° North; **Supplementary Figure S1**). Subsequently we used the transfer function provided by Lehnert et al. (2018) to relate hydrogen isotope ratios in fur (δ^2^H_f_) values as a linear function to δ^2^H_P_ values. This enabled us to directly map the assignment samples onto the isoscape: we used the *isofind* function of the package to test the probability of a sample originating from the candidate location Prieros, Brandenburg, Germany, under the null hypothesis that the unknown location of origin is identical to the candidate location. For assignments where the p-value fell below α = 0.05, we rejected the null hypothesis and concluded a migratory status for the respective individual. Finally, assignments were visualized highlighting areas with the most probable place of origin (**Figure 2**). Additionally, we tested whether the location of origin for a particular fur sample could be predicted by a triple isotope assay, using a discriminant function analysis (DFA) of the *“MASS*” package: DFA uses linear combinations of the predictor variables (here: δ^2^H, δ^13^C and δ^15^N) to classify a response variable (here: migratory or local status). We used 70% of the dataset as training set and the remaining 30% as testing set and then fitted linear discriminant analysis (LDA) models to both data sets. The migration strategy assignment (long-distance migrant or local) received 95% reliability support using a triple isotope approach, which is in line with previous reports using this method (93% reliability support in: Popa-Lisseanu et al., 2012).

**Figure 2.**
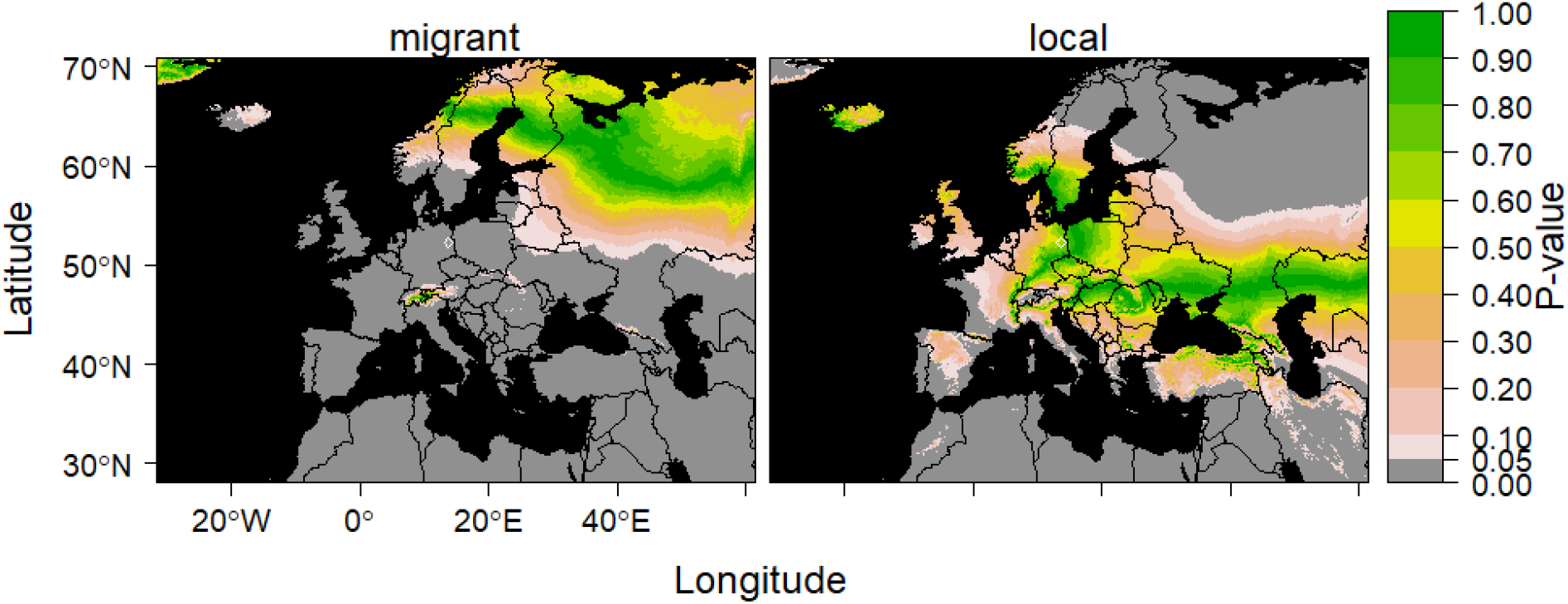
Geographic probability distribution of the most probable place of origin of a) an assigned migrant and b) an assigned local bat. Colour scale indicates level of certainty of correct assignment (expressed as p-level). Bats for which a local status could be excluded using an α = 5% threshold, were assigned a migrant status. Bats for which a local origin could not be excluded were assigned a local status, i.e. originating from the sampling location Prieros, Brandenburg, Germany (∼40 km south-east from Berlin, indicated on the maps by the white diamond).

#### Emergence, spatial activity and acoustic exploration behaviour

Following Schabacker et al. (2021) we focused on three relevant and independent response measures: full body emergence (yes/no), total number of chambers visited, and acoustic exploration. Latency to full body emergence correlated strongly with the two other emergence response measures (latency to head emergence: r_s_ = 0.97, P < 0.001 and emergence duration: r_s_ = 0.41, P = 0.005), and the total number of chambers visited correlated strongly to the other spatial activity response measure, the number of unique chambers visited (r_s_ = 0.77, P < 0.001). Full body emergence and the total number of chambers visited were therefore used as representative measures for emergence and spatial activity behaviour, respectively.

To test if response to a novel environment varies with migration strategy, three logistic regression models were constructed, with migration strategy (local/migrant) as dependent variable and (A) body emergence (yes/no), (B) total number of chambers visited, or (C) acoustic exploration as independent variable of interests. We conducted backwards stepwise model selection always keeping the independent variables of interest in the model. Control variables included the season the sample was taken (April or October), since occurrence of migrants is seasonal (Lehnert et al., 2014), the body condition of the subject as weight (g) divided by length of the lower arm (mm), and the length of the lower arm (mm), both which were previously shown to predict migration strategy (Lehnert et al., 2018). Control variables with P ≥ 0.1 were removed from the model. For eight assays, the migration strategy could not be reliably determined (amplitude peak of the δ^2^H sample too high), resulting in a final sample size of (A) N = 82, (B) N = 41, (C) N = 41 for first assays. Due to the low number of repeated assays (N = 9), we decided to focus our analysis on the behavioural responses to the first assay only. Model assumptions were verified using the packages ‘dHARMA’ (Hartig & Hartig, 2017) and ‘performance’ (Lüdecke et al., 2021), using the functions: check_collinearity(), simulateResiduals() and plot(). The data and code underlying the analyses can be accessed here: https://osf.io/kxtj5/?view_only=bdb1ecb24bf1400d9a19152c29c133cc

## Results

Based on the stable isotope analyses, 60% (49 out of 82 valid samples) of the bats came from a distant origin (**Supplementary Table 1, Supplementary Figure S2**. When exposed to the novel environment for the first time, only 51% of the bats (45 out of 89) emerged and these were most likely to be locals (glm: Estimate (SE) = −1.01 (0.48), Chisq = 4.43, N = 82, P = 0.04, **Figure 3a**). We rule out the possibility that differences in handling experience influenced the outcome of this study, because distant migrants were as likely to be previously handled (banded) as local conspecifics (Fisher Exact Test: OR (95% CI) = 0.57 (0.20 - 1.54), N = 89, P = 0.26).

**Figure 3.**
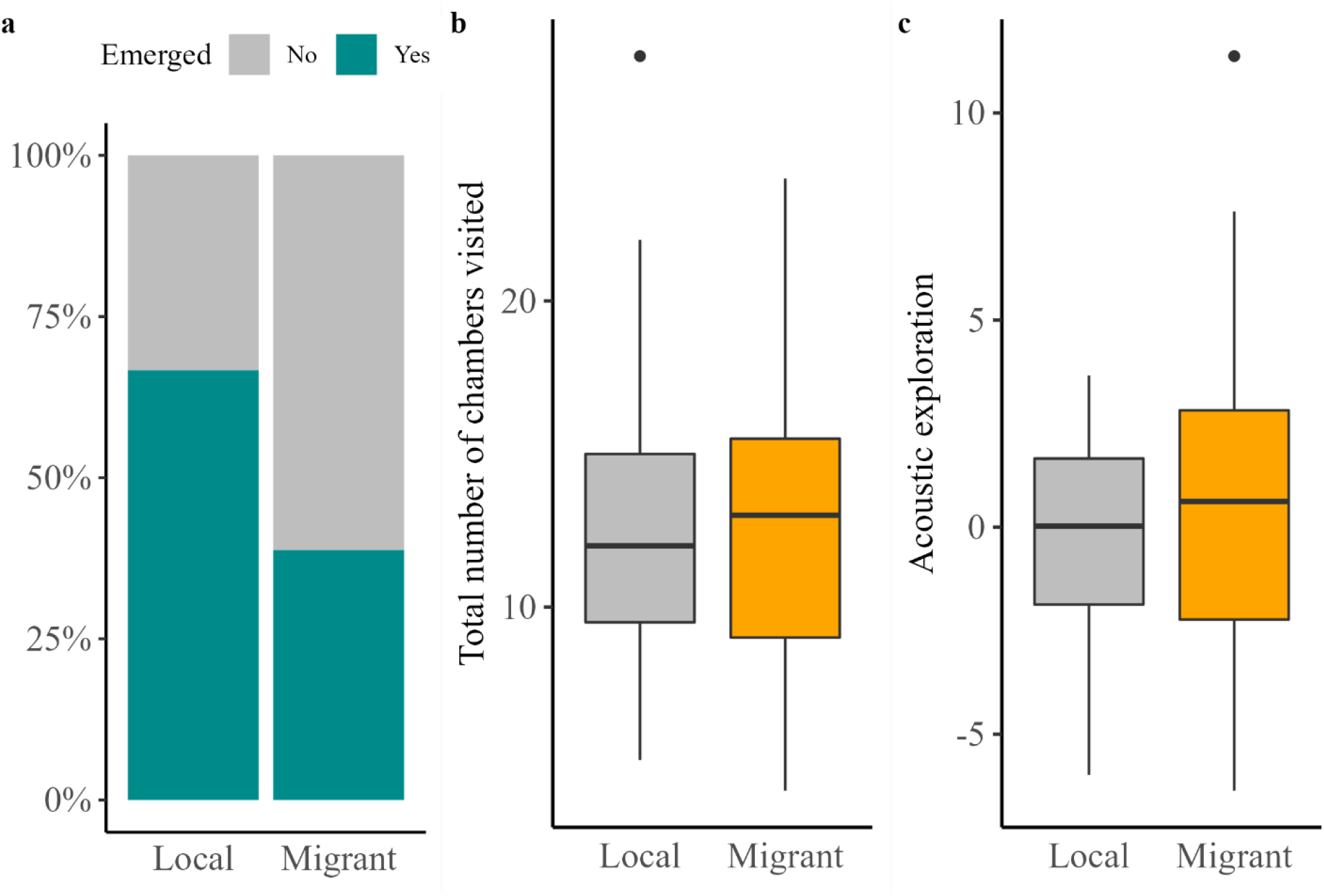
Noctule bats behavioural responses to the novel environment assay as function of their migration strategy. Local individuals were (a) more likely to emerge into the novel environment (maze), yet local individuals and long-distance migrants did not differ in (b) their spatial activity (total number of chambers visited) or (c) their acoustic exploration (residuals of the number of echolocation calls emitted over the number of chambers visited).

Individuals that emerged into the novel environment showed substantial variation in their response (**Supplementary Figure S3a-e**). As expected, the number of calls emitted after emergence during the first test was strongly correlated to the total number of chambers visited (both variables square root transformed; r = 0.42, N = 45, P = 0.004; **Supplementary Figure S4a**). Despite this strong correlation, there was substantial variation in how much bats under or oversampled the novel environment (i.e., acoustic exploration, **Supplementary Figure 4b**). Nevertheless, migrants were not characterized by higher spatial activity (the number of chambers visited) (glm: Estimate (SE) = −0.02 (0.06), Chisq = 0.14, N = 41, P = 0.70**; Figure 3b**) nor stronger acoustic exploration in the novel environment (glm: Estimate (SE) = 0.11 (0.09), Chisq = 1.33, N = 41, P = 0.25; **Figure 3c**).

There were no notable correlations between the three behavioural responses of interest themselves (i.e., emergence latency, number of chambers visited, and acoustic exploration) nor between these three behavioural responses and the number of air puffs and number of head pokes into adjacent chambers (**Supplementary Table S2**).

## Discussion

The environment is changing and so are the migration strategies of species worldwide. Yet little is known about how migrants and locals of the same species differ in how they cope with novel environments. Using a unique combination of isotopic geolocation and an in-situ novel environment assay, we unexpectedly found, that local individuals were more likely than distance migrants to emerge into the novel environment. These findings suggest that local bats may more pro-actively cope with novelty than migrants. This could make it easier for local bats to adjust to human-induced changes but may also put them at more risk if these changes include danger. In contrast, there were no correlations between migration strategy and spatial activity or acoustic exploration, suggesting that explorative behaviour, once past the initial hurdle of emergence or approach, is equally important for migratory and local noctule bats.

### Local individuals are more likely to enter novel environments

We investigated if bats with different migration strategies are characterized by different responses to a novel environment. For this, we quantified the emergence into a novel environment, the spatial activity in a novel environment, and acoustic exploration (Schabacker et al., 2021). We found a correlation between emergence behaviour and migration strategy, but in the opposite direction than expected. As migrants are more often confronted with novel environments, we expected migrants to be more likely to emerge into the novel environment. Yet our study shows that the bats that emerged were more likely to be local individuals than long-distance migrants. This appears to contrast with related studies in other taxa. For example, roach (*Rutilus rutilus*) that emerged quicker from a refuge migrated more often (Chapman et al., 2011). In bank voles (*Myodes glareolus*), individuals with shorter latencies to emerge and investigate an unknown area occupied larger home ranges (Schirmer et al., 2019). Similarly, in golden-mantled ground squirrels (*Callospermophilus lateralis*), individuals that fled later when approached by a human observer maintained larger core areas, and exhibited elevated movement speeds (Aliperti & Todgham, 2021). In two warbler species (*Sylvia spec*.), more individuals from the migratory species entered a novel environment than individuals from the closely related local species, and they did so more quickly (Mettke-Hofmann et al., 2009; Mettke-Hofmann & Greenberg, 2005).

Fast emergence into a novel environment from a secure place can be interpreted as a form of risk-taking (Carter et al., 2013). As local individuals spend extended periods of time in the same location, they might have more complete information about their current local environment, simply because they had more opportunities to sample this environment (Dall et al., 2015). These individuals could then be expected to be very well informed about predation risks throughout the year, and thus might behave more confidently, i.e., risk prone (Error management theory: Johnson et al., 2013; Feyten et al., 2019). This could facilitate quick adaptation to novel changes to the environment, but may also expose local bats to more to danger (e.g., wind turbines or unsuitable roosts). On the other hand, migratory individuals are more often confronted with novel environments, where risk-assessment is difficult and dangerous. Migrants might benefit from behaving warily and vigilantly, because they do not have extended knowledge about predation danger in their current environment (Dangerous niche hypothesis: Greenberg & Mettke-Hoffmann, 2001; Mettke-Hofmann et al., 2013). A more risk-averse behaviour in migrants could thus compensate for the increased risks associated with migration behaviour, but could also imply that migrants would need more time to adjust to novelty (e.g., uptake of artificial bat boxes) and suffer more stress when confronted with novelty.

Exposure to novelty increases effects associated with physiological stress (Pfister, 1979), yet not all individuals are affected equally. Migratory bats, which were less eager to enter a novel environment, are likely more affected by novelty-associated stress. Studies with free-ranging sparrows showed that when confronted with novelty, inquisitive birds exhibited the lowest stress response, measured as stress-induced corticosterone (Lendvai et al., 2011). This has important implications, since energy attributed to stress responses will not be available for other energetically demanding activities, such as migration. Giving the rapidly changing world, especially for migratory bats which are continuously confronted with anthropogenically changed roosts and stopover sites (Voigt & Kingston, 2016), future research should investigate if migrants and locals indeed differ in their endocrinological stress responses to novelty. Animal migration is under pressure (Wilcove & Wikelski, 2008) and a better understanding of the underlying mechanism that may make migrants more vulnerable to changing environments would allow for more targeted conservation plans.

### No difference in spatial activity and acoustic exploration between migration strategies

Bats actively manipulate the level of echolocation pulses they emit, and thus individually control incoming information from the environment. We expected migrants to show more acoustic exploration as they could benefit more from continuously updating their information when moving through novel environments. When migrants and locals differ in their ability to detect changes in the environment, one could have more difficulty locating suitable novel roosting spots or avoiding fatal collisions with novel anthropogenic objects (e.g. wind turbines: Lehnert et al., 2014; Voigt et al., 2012) than the other, resulting in selective disappearance of a migration strategy. This is relevant since noctule bats are a common victim of wind turbines (Kruscynski et al. 2022, Lehnert et al., 2014). However, despite substantial variation in spatial activity and acoustic exploration between tests, we found that locals and migrants did not differ in these explorative behaviours. Our results thus suggest that spatial activity and acoustic exploration is equally used by locals and migrants, possibly because these behaviours are also crucial outside the migration context, such as during foraging and orientation inside the roost (Kunz, 1982; Schnitzler et al., 2003; Thomas et al., 2004). In our assay, we let bats choose whether they entered the novel environment, since forcing non-emerging bats into the novel environment would result in primarily fear-related behavioural responses rather than exploration. We, however, acknowledge that with only ca. 50% of the bats emerging, it is possible that migrants that did emerge were not representative of the general level of acoustic exploration and spatial activity exhibited by migrants.

If acoustic exploration is indeed not linked to migration strategy, it remains an open question what else could explain the substantial variation in acoustic exploration we found. In an earlier study on another tree-roosting species, the Nathusius bat, we found both spatial activity and acoustic exploration to consistently differ between individuals (Schabacker et al., 2021). Unfortunately, we were not able to collect enough repeated measures to test repeatability in noctule bats, as recapture success was too low (only nine individuals). We do see interesting parallels with the Nathusius bats, as they also showed substantial levels of variation in spatial and acoustic traits and a strong correlation between spatial activity and echolocation activity. In contrast, we did not find acoustic exploration in noctule bats to vary with other exploratory behaviours, such as number of head pokes into adjacent chambers, as we found in Nathusius’ bats (Schabacker et al., 2021). We can thus not draw conclusions about whether the acoustic exploration we measured in noctule bats is representative for an exploration-related personality trait. We hope to instigate future research focusing on the repeated examination of acoustic exploration in echolocating bats. This could be achieved by housing captured bats temporarily on site, rather than depending on recapture, and conducting repeated tests, prior to release. For our current study, the first of its kind in bats, we opted for the least disruptive approach and so maximize ecological validity.

### Isotopic geolocation to study migration ecology in an understudied taxon

Traditionally, migration behaviour is studied by means of extrinsic markers, which have facilitated crucial insights into animal movement (Hobson, 2018; Rutz & Hays, 2009), but also have their limitations (Robinson et al., 2010). This is especially so when working with elusive long-distance migrants with small body weights (Kays et al., 2015). Radio transmitters only allow for short to medium distance tracking and often require extraordinary effort. Most geolocators must be recaptured to retrieve data and satellite telemetry devices are often too large for many migratory animals (Kays et al., 2015; Nathan et al., 2022). Lastly, external markers may alter an individual’s (movement) behaviour. Here, we circumvented these limitations by using endogenous stable isotope ratios as intrinsic markers in non-invasive isotopic geolocation (Popa-Lisseanu et al., 2012). Using stable hydrogen isotope analysis, we were able to assign migration strategies to 82 individual female common noctule bats.

Stable isotope analysis has, nevertheless, some limitations. Firstly, in this study, we used a transfer function recently established for common noctule bats (Lehnert et al., 2018). This transfer function uses a combination of δ^2^H values of local, non-migrating insectivorous bat species and those of noctule bats sampled during their non-migratory period across Central and Eastern Europe (Lehnert et al., 2018; Voigt et al., 2014; Voigt & Lehnert, 2018). Transfer functions are crucial to correctly relate δ^2^H ratios of precipitation water to that of bat fur (Voigt & Lehnert, 2018). Such transfer functions work best if they are taxon- or guild specific, as food related differences in δ^2^H ratios across species could cause differences in their respective isotopic composition, albeit animals being exposed to the same δ^2^H values of precipitation and ground water (Voigt et al., 2015). However, to date, we generally lack data to create species-specific transfer functions (Voigt & Lehnert, 2018). If available, single species transfer functions often suffer from a small sample size and thus might not cover the full range of variation in the δ^2^H tissue values. As only a relatively small sample size would have been available for a common noctule bat specific transfer function, we opted for a multispecies-informed transfer function, thus resulting in a lower spatial resolution but generally a more comprehensive and reliable assignment (Voigt & Lehnert, 2018). Secondly, the isoscapes generated with isotopic geolocation depend on the input of precipitation data and are thus limited in its predictive power in regions sparsely covered with weather stations. Additionally, regions of similar δ^2^H_P_ values naturally yield a similar probability of origin for the respective assignments. This is especially true in the northern hemisphere, where isoclines (regions of similar δ^2^H_P_ values) follow latitude closely and thus resolve animal movements from east to west poorly (Voigt & Lehnert, 2018). However, in this study, we focused on the differentiation between local individuals and long-distance migrants, whereby the exact place of origin was not determinative, but the assigned distance from the sampling point. Our study thus demonstrates the potential of isotopic geolocation as a powerful tool to assess cryptic migratory behaviours of elusive taxa, such as bats.

## Conclusion

Our unique approach, combining behavioural assays with isotopic geolocation in an elusive and vulnerable taxon, gave us unexpected insights, thereby highlighting the importance of studying the behavioural correlates of migration across various distinct taxa. Contrasting to our hypotheses, we found local individuals to be more likely to enter an unfamiliar environment than long-distance migrants and we did not detect a relationship between spatial activity or acoustic exploration and migration strategy. This appears in contrast to similar studies on behavioural trait variation and (long-distance) movement strategies in other taxa. Revealing such taxon-specific relationships between migration strategies and other behavioural traits will be essential for accurately predicting within-species vulnerability to environmental change and for formulating effective conservation plans.

## Acknowledgements

We thank Kseniia Kravchenko for assistance in the isotope analysis, Anja Luckner for the conduction of the laboratory analyses and Ana Paul for support during fieldwork. L.S. was funded by a Humboldt Research Fellowship for Postdoctoral Researchers (Ref 3.3 – NLD – 1192631 - HFST-P) awarded by the Alexander von Humboldt-Stiftung. We thank Rebecca Scheibke for her illustration of the novel environment assay.

## Supplementary information

### Tables

**Table S1.**
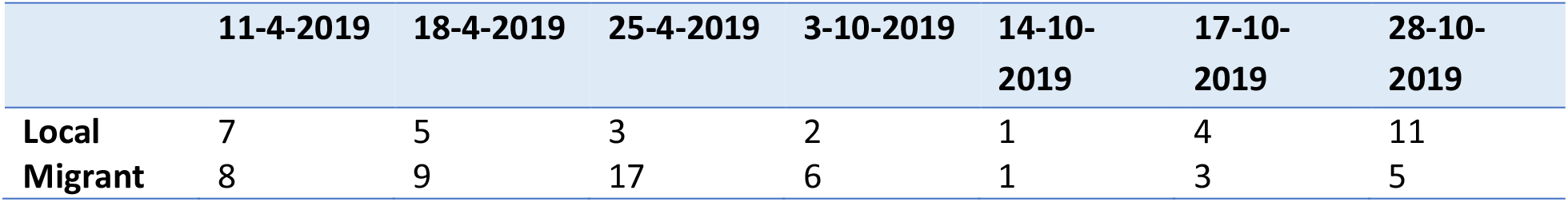
Migration strategy assignment across first test days (N = 82, due to 7 invalid readings).

|  | 11-4-2019 | 18-4-2019 | 25-4-2019 | 3-10-2019 | 14-10-2019 | 17-10-2019 | 28-10-2019 |
| --- | --- | --- | --- | --- | --- | --- | --- |
| **Local** | 7 | 5 | 3 | 2 | 1 | 4 | 11 |
| **Migrant** | 8 | 9 | 17 | 6 | 1 | 3 | 5 |

**Table S2.** Correlations between the behavioural responses of interest and additionally recorded behavioural responses.

| Response 1 | Response 2 | $r_s$ | P | P_adjust (Holm) |
| --- | --- | --- | --- | --- |
| **Emergence latency (s)** | Total number of chambers visited | -0.11 | 0.47 | 1.00 |
|  | Number of air puffs | -0.13 | 0.39 | 1.00 |
|  | Number of head pokes | -0.21 | 0.16 | 1.00 |
|  | Acoustic exploration | -0.12 | 0.42 | 1.00 |
| **Total number of chambers visited** | Number of air puffs | -0.32 | **0.03** | 0.27 |
|  | Number of head pokes | -0.14 | 0.36 | 1.00 |
| **Acoustic exploration** | Number of air puffs | -0.06 | 0.69 | 1.00 |
|  | Number of head pokes | -0.09 | 0.57 | 1.00 |

### Figures

**Figure S1.**
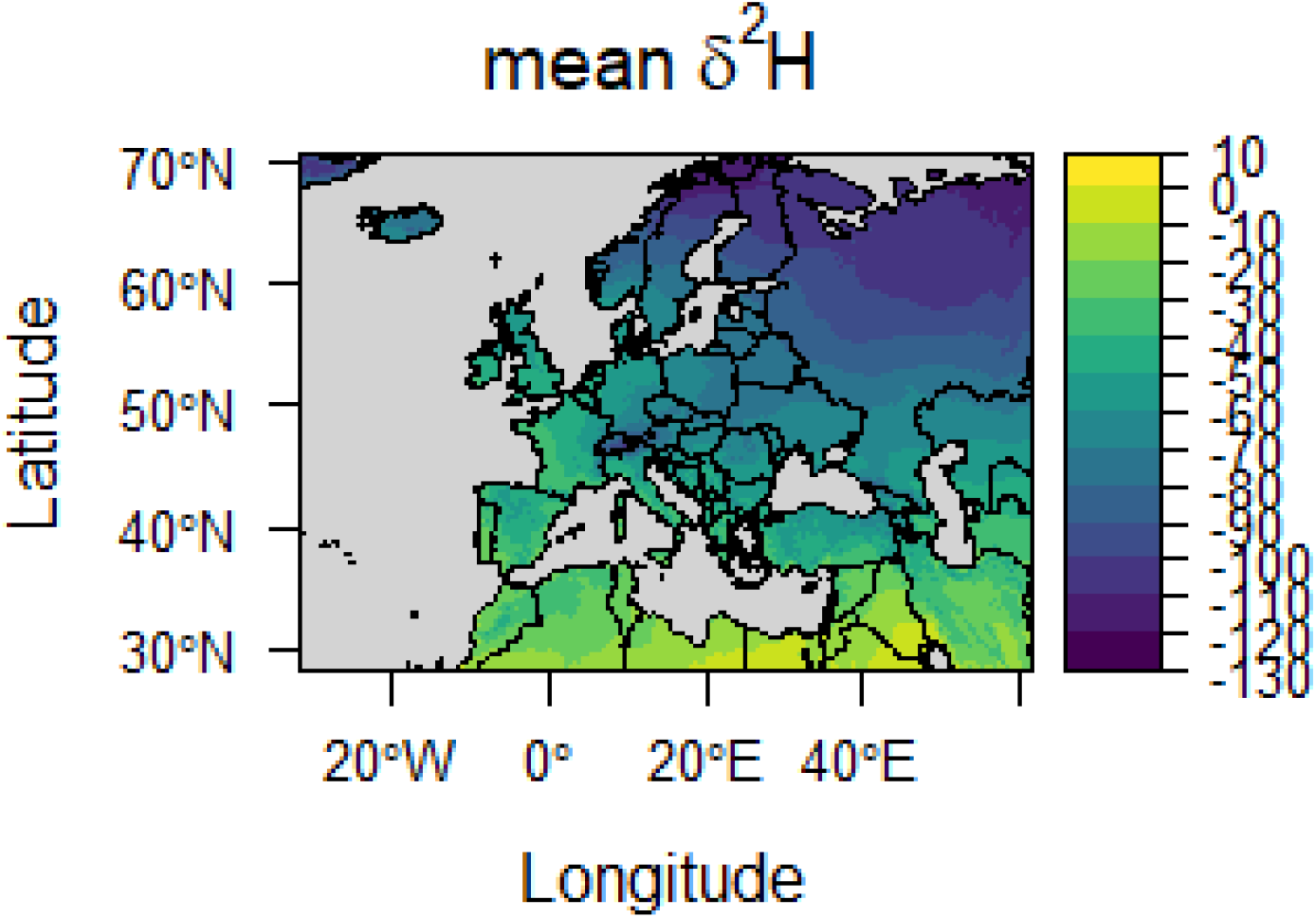
Europe isoscape. Graphic visualization of mean δ^2^HP values roughly covering Europe (longitude range: −30° East −60° West, latitude range: 30 North° – 70° North)

**Figure S2.**
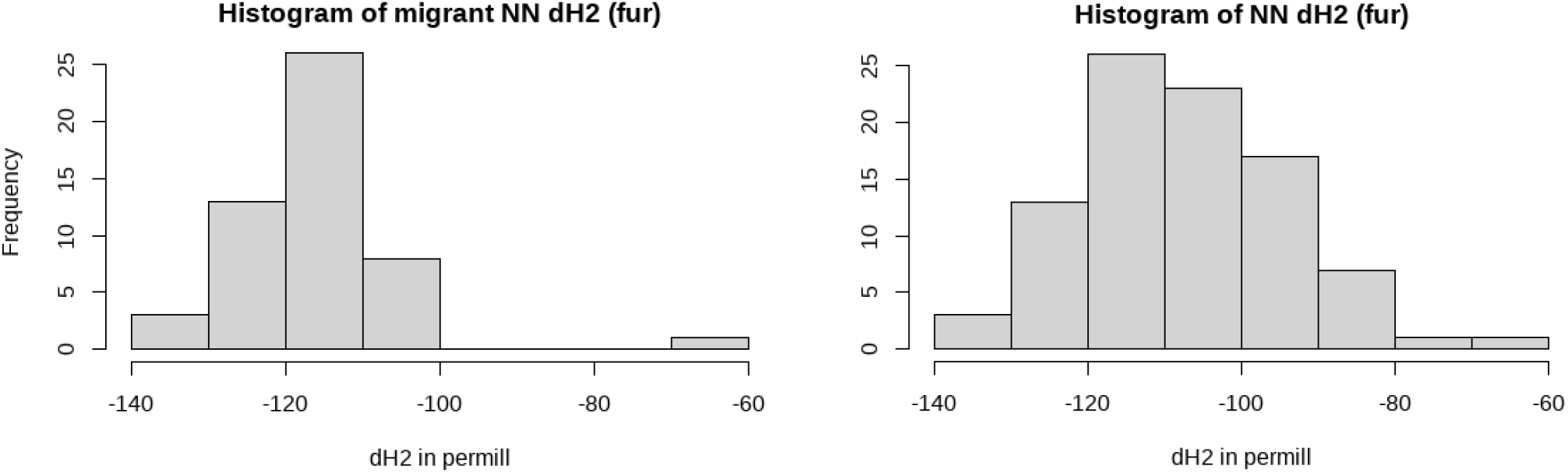
Histograms of δ^2^H fur values of a) migratory and b) local *Nyctalus noctula*

**Figure S3.**
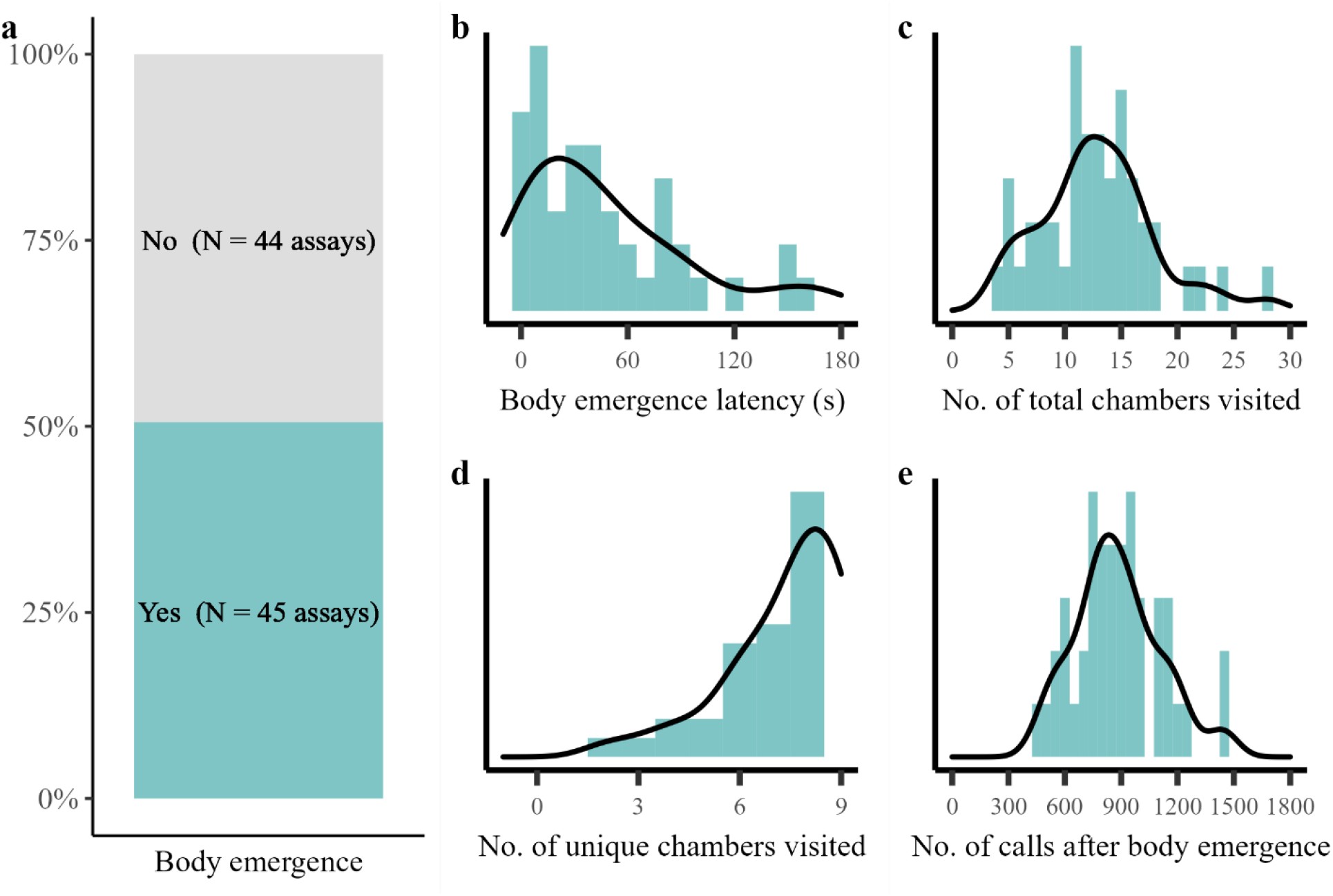
Behavioural responses of common noctule bats to the novel environment assay. (a) Half of 90 female bats emerged into the novel environment (maze). The bats showed ample variation in (b) latency to emerge, (c) total number of chambers visited, (d) unique number of chambers visited (nine unique chambers were available), and (e) the number of echolocation calls emitted in the maze.

**Figure S4.**
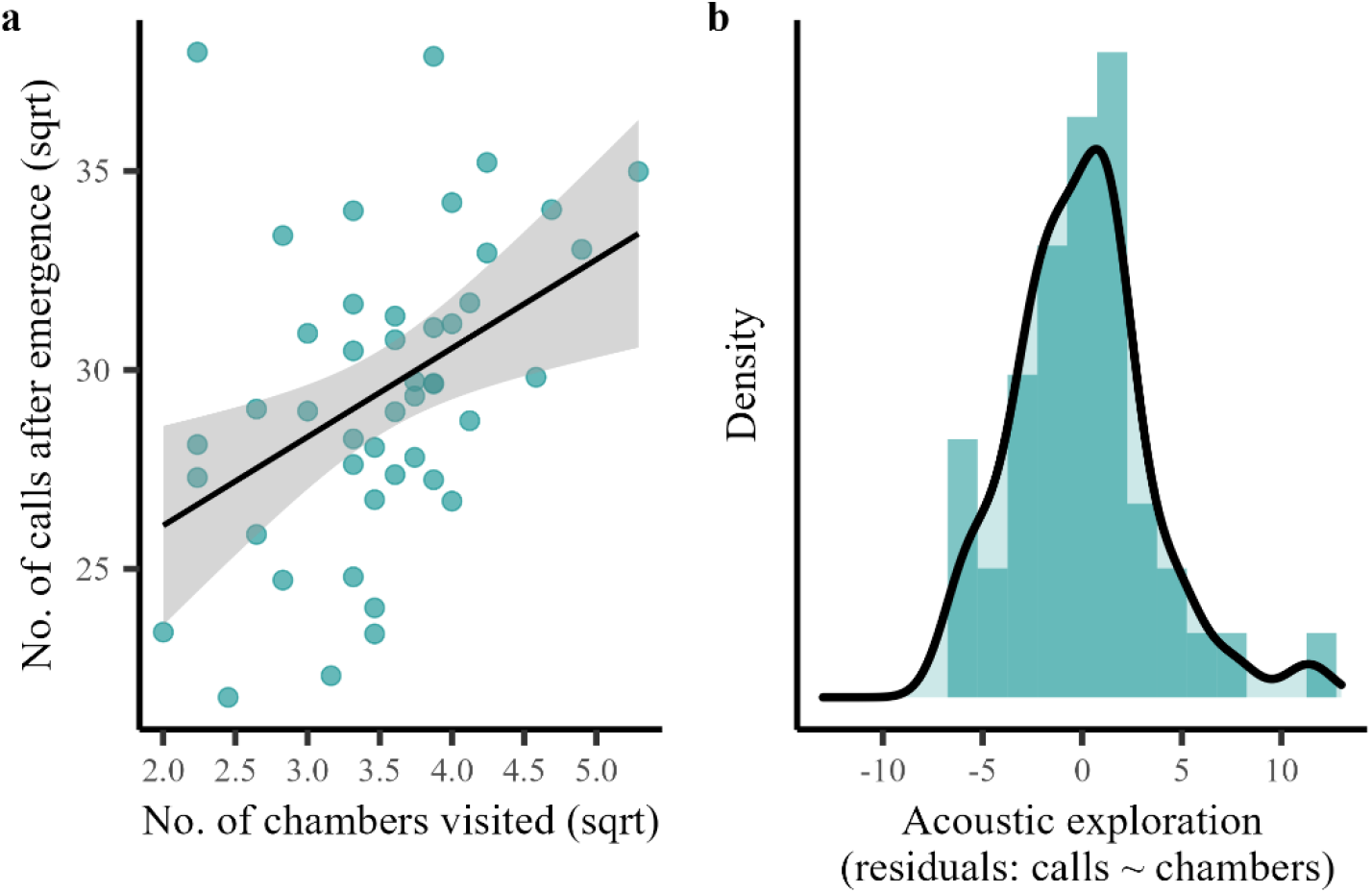
Number of echolocation calls emitted in the novel environment as function of the number of chambers an individual bat visited. (a) The number of echolocation calls was positively correlated with the number of chambers visited during the assay. (b) There was substantial variation in the correlation between the number of chambers visited and the number of echolocation calls emitted, indicating bats that were under or oversampling their environment, visualized by the histogram of the residuals of this correlation.

